# A non-canonical role for UPR^ER^ during heat stress in *C. elegans*

**DOI:** 10.64898/2026.01.08.698479

**Authors:** Athena Alcala, Toni Castro Torres, Rebecca Aviles Barahona, Phillip A Frankino, Ryo Higuchi-Sanabria, Gilberto Garcia

## Abstract

Organisms rely on coordinated stress responses to maintain cellular homeostasis. Perhaps the best-known example of multiple stress inputs converging onto a single response is the integrated stress response (ISR), which reduces global translation under various stress conditions to reduce the protein folding burden of the cell. Similarly, most stress responses generally involve coordination of additional protein homeostasis (proteostasis) pathways, including increased expression of chaperones to refold proteins, as well as activation of clearance mechanisms, such as autophagy and the ubiquitin proteosome system. Our study investigates how heat stress can influence coordinated activation of both cytosolic and ER chaperones, exploring bidirectional cross talk between canonical activators of the cytosolic heat-shock response (HSR) and the unfolded protein response of the ER (UPR^ER^). Using robust transcriptional reporters in the *C. elegans* model system, we explore a non-canonical activation of the UPR^ER^ under heat stress by the coordinated effects of XBP-1 and HSF-1. We further investigate inter tissue communications of stress whereby neuronal or glial activation of the UPR^ER^ can result in heterotypic enhancement of the HSR in peripheral and can increase thermotolerance. This work highlights the complex convergence of cellular stress responses, a phenomenon that may reflect a general strategy wherein localized stress can activate numerous proteostasis pathways to prevent whole cell and whole organism damage.

**Article Summary:** A reductionist approach to studying cellular stress responses is critical for dissecting specific molecular and genetic drivers of stress response. However, stress responses are often convergent and overlapping, and these single input and output studies may miss their complex interplay. Many studies have revealed the intricate coordination of stress responses, including the ability of seemingly organelle-specific stress responses, like mitochondrial stress responses, to directly influence cytosolic and ER health. Our study adds to this growing field by describing a unique, bidirectional crosstalk between the cytosolic and ER stress pathways, highlighting systemic coordination of stress resilience.

## Introduction

Organisms are constantly exposed to stress, which can have detrimental effects on health and physiology. As such, several regulatory stress responses function to promote and maintain cellular homeostasis, tailored for each specific organelle. For example, the unfolded protein response of the endoplasmic reticulum (UPR^ER^) is activated under ER stress by unconventional splicing of *xbp-1* mRNA by the ER-resident transmembrane protein, IRE-1. This results in translation of functional XBP-1s to activate transcription of genes essential for ER function (Walter and Ron 2011; Frakes and Dillin 2017). In addition to this most well) anded IRE-1/XBP-1 arm, two additional branches of UPR^ER^ are mediated by protein kinase R-like endoplasmic reticulum kinase (PERK, or PEK-1 in *C. elegans*) and activating transcription factor 6 (ATF-6). Similar to IRE-1, PEK-1 and ATF-6 are ER-resident transmembrane proteins containing a luminal domain that can bind unfolded proteins in the ER (Schröder and Kaufman 2005). Under ER stress, PEK-1 activation results in phosphorylation of eukaryotic initiation factor (EIF-2A) to reduce global translation and reduce the protein folding burden of the ER (Ma and Hendershot 2003; B’chir et al. 2013; Han et al. 2013). ATF-6 activation involves cleavage of the cytoplasmic domain and translocation into the Golgi for further processing (Haze et al. 1999; Ye et al. 2000).

A similar machinery, the heat-shock response (HSR), exists for the cytosol, which is regulated by heat-shock factor 1, HSF-1. Under normal, non-stressed conditions, HSF-1 is maintained in an inactive form by direct binding with HSP-70 and HSP-90, which act as negative regulators of HSF-1 function (Shi et al. 1998; Anckar and Sistonen 2011; Neef et al. 2014). In response to stress, these molecular chaperones are titrated away from HSF-1, allowing for a series of post-translational modifications that promote trimerization and nuclear translocation of HSF-1, allowing for transcriptional activation of HSR targets (Anckar and Sistonen 2011; Dai 2018).

Much of the foundational research in stress biology has understandably used a reductionist approach studying individual stress response pathways within specific organelles, enabling careful dissection of their molecular mechanisms. However, growing evidence suggests that stress responses often have pleiotropic effects, influencing multiple organelles and systems, highlighting the need to also consider inter-organelle communication and systemic coordination of stress. For example, low levels of mitochondrial stress can increase functional capacity of the HSR, delaying age-associated loss of HSR function to promote stress resilience and longevity (Labbadia et al. 2017). In addition, knockdown of *hsp-6*, which encodes a mitochondrial chaperone, resulted in activation of the cytosolic HSR through a complex signaling cascade involving fatty acid remodeling (Kim et al. 2016). Finally, HSF-1 impacts mtDNA gene expression by elevating histone H4 levels and altering mtDNA chromatin state, which can activate the unfolded protein response of the mitochondria (UPR^MT^) to promote longevity (Sural et al.).

Similar overlapping functions of the UPR^ER^ and HSR have also been observed. One study found nine stress responsive genes overlapping under ER-stress and heat-stress conditions (Liu and Chang 2008), including the ER chaperone, HSP-4/BiP (Kohno et al. 1993) and the COPII cargo receptor, Erv29 (Caldwell et al. 2001). In addition, activating the HSR through a constitutively active form of HSF-1 has been shown to compensate for growth defects in cells lacking a functional UPR^ER^. Heat-shock has also been shown in mammalian cells to activate canonical XBP1 targets (Heldens et al. 2011). These results are not surprising considering the pleiotropic effects of heat stress exposure: exposure to heat would likely result in protein unfolding in all cellular compartments and not just the cytosol, thus likely requiring activation of multiple stress response pathways. However, whether heat-shock can directly activate UPR^ER^ in *C. elegans*, and what specific stress response pathways are required for this activation have yet to be studied. One study demonstrated that *C. elegans* electrotaxis behavior is regulated by interconnected cytosolic, mitochondrial, and ER stress response pathways, including the HSR and UPR, further suggesting an overlapping and coordinated nature of these stress responses. Therefore, in this study, we directly tested the impact of exposure to heat-stress on UPR^ER^ activation, and identified a non-canonical HSF-1 dependent UPR^ER^ in response to heat. In addition, we found that neural overexpression of *xbp-1s* previously found to promote whole-organism UPR^ER^ activation to extend lifespan (Taylor and Dillin 2013; Frakes et al. 2020) can similarly boost the HSR to promote thermotolerance to a comparable level as neuronal *hsf-1* overexpression (Douglas et al. 2015). Overall, our study adds further evidence to the complex interplay and general overlap between cellular stress signaling networks.

## Materials and Methods

### *C. elegans* maintenance

All strains used in this study are derived from N2 wild-type worms and were previously constructed, validated, and published in previous studies. All reporter strains are available on Caenorhabditis Genetics Center (CGC) as referenced in reagents, and transgenic animal strains will be made available upon request. All animals are maintained at 15°C fed OP50 *E. coli* B strain during maintenance. For experiments, animals are synchronized using a standard bleaching protocol as described previously (Torres et al. 2022). Briefly, worms are collected into a 15 mL conical tube using M9 solution (22 mM KH_2_PO_4_ monobasic, 42.3 mM NaHPO_4_, 85.6 mM NaCl, 1 mM MgSO_4_) and subjected to a bleaching solution (1.8% sodium hypochlorite, 0.375 M NaOH in M9) at a 20:1 bleach:worm volume ratio. Bleach:worm mixture is vigorously shaken until all warm carcasses are dissolved, and intact eggs are washed four times using M9 solution by centrifugation at 1,100 x g for 1 min. After the final wash, animals were L1 arrested by incubating overnight in 12 mL M9 solution at 20°C on a rotator for a maximum of 24 hours. After bleaching, worms were then plated onto HT115 *E. coli* K strain from L1 until day 1 adulthood where all experiments were performed.

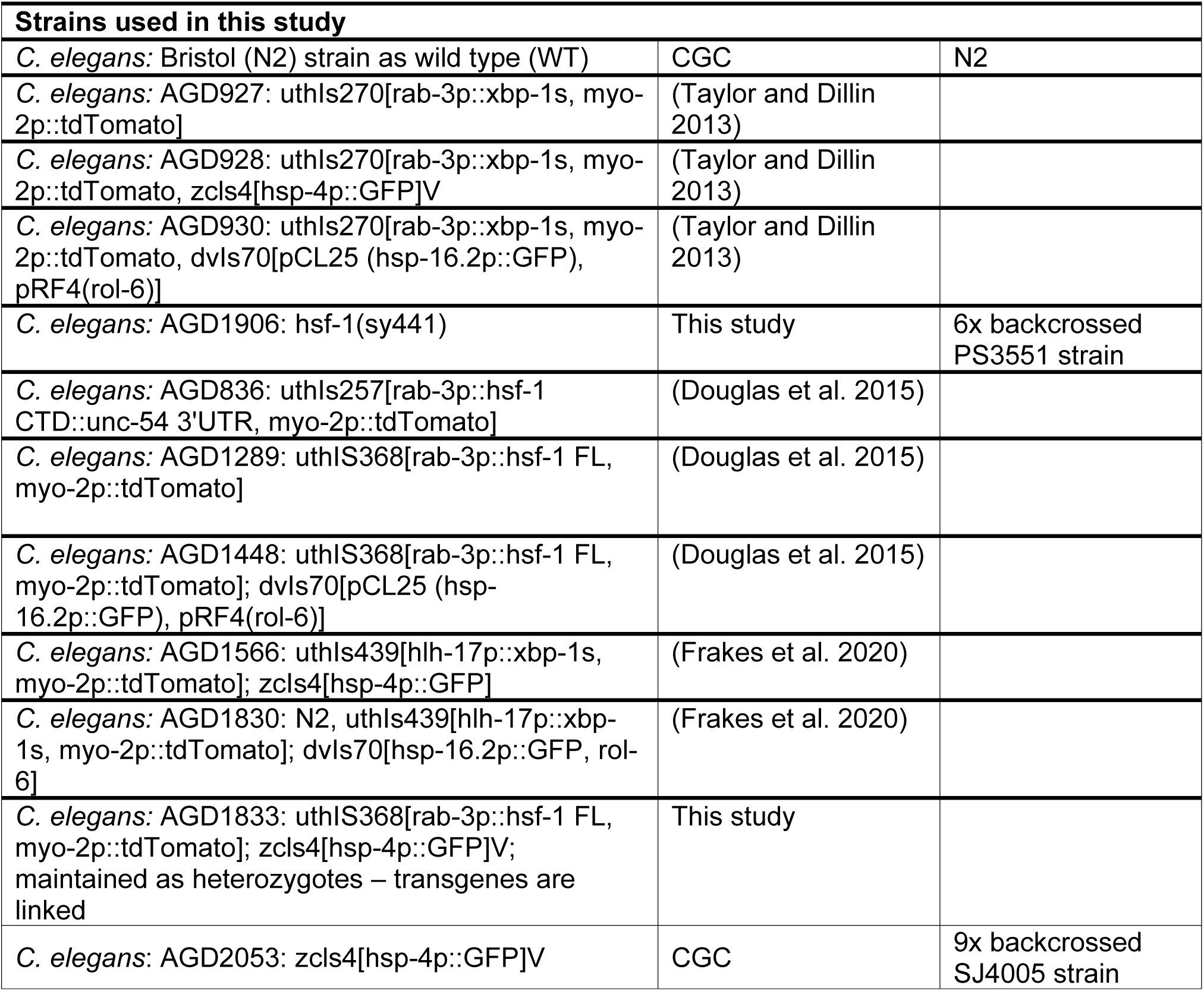
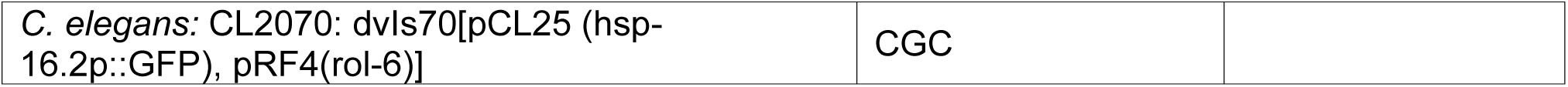

### Plates

Standard NGM plates for maintenance using OP50 contained the following: Bacto-Agar (Difco) 2% w/v, Bacto Peptone 0.25% w/v, NaCl_2_ 0.3% w/v, 1 mM CaCl_2_, 5 µg/ml cholesterol, 0.625 mM KPO_4_ pH 6.0, 1 mM MgSO_4_.

Standard NGM-RNAi plates for experimental condition using HT115 bacteria contained the following: 2% RNAi plates for experiments contained the following: Bacto-Agar (Difco) 2% w/v, Bacto Peptone 0.25% w/v, NaCl_2_ 0.3% w/v, 1 mM CaCl_2_, 5 µg/ml cholesterol, 0.625 mM KPO_4_ pH 6.0, 1 mM MgSO_4_, 100 µg/mL carbenicillin, 1 mM IPTG.

### *C. elegans* microscopy

All imaging of reporter strains were performed as previously described (Bar-Ziv et al. 2020) at day 1 of adulthood using the synchronization method described above. For all RNAi experiments, animals were exposed to RNAi starting from the L1 stage. Specifically, the heat-shock reporter *hsp-16.2p::GFP* was imaged by performing heat-shock at 34°C for 2 hours in a heated incubator, followed by recovery at 20°C for 2 hours. Control animals were left at 20°C during the duration of the experiment. For the UPR^ER^ *hsp-4p::GFP* and *hsp-6p::GFP* experiments, animals are heat-shocked at 34°C for 2 hours in a heated incubator, followed by recovery for 4 hours.

For tunicamycin experiments, L4 animals are washed off plates and exposed to 25 ng/µL tunicamycin or 0.1% DMSO (control) for 4 hours in M9 solution in a rotator at 20 °C. Tunicamycin or DMSO are washed 3x with M9 and animals are recovered on standard HT115 plates overnight (up to 16 hours) and imaged at day 1 of adulthood (Bar-Ziv et al. 2020).

All imaging is performed by picking 10+ animals into a 5 μL pool of 100 mM sodium azide on a standard NGM plate without bacteria to induce paralysis. Once the sodium azide droplet dried, paralyzed animals were then lined up alongside each other and imaged under a Leica M205FCA automated fluorescent stereomicroscope running LAS X software and equipped with a standard GFP filter, Leica LED3 light source, and Leica K5 camera. For all imaging experiments, 3 biological replicates were performed with 2 technical replicates each, and 1 representative image was chosen for use in figures. For quantification, Fiji (Schindelin et al. 2012) was used to draw a region of interest along each individual worm, and integrated density was measured and normalized to area.. Graphing and statistical analysis were performed with GraphPad Prism 10 software using a Mann-Whitney test or Kruskal-Wallis testing for multiple comparisons unless otherwise stated.

### Thermotolerance measurements

For thermotolerance experiments, animals were grown on RNAi plates on either EV or RNAi bacteria from L1 stage. Day 1 adult animals were placed at 34°C and animals were scored for viability every 2-3 hours until all animals were scored (Torres et al. 2022). All assays were performed on >3 biological replicates. Survival curves were plotted using GraphPad Prism 10 software and statistics were performed using a Log-Rank test in GraphPad Prism 10.

### RNA-sequencing

RNA collection was performed on ∼800 animals at day 1 of adulthood. For heat-shock, animals were heat-shocked at 34°C for 2 hours and allowed to recover at 20°C for 1 hour prior to sample collection. Control animals were continuously grown at 20°C until sample collection. Animals were collected into Trizol solution and worms were freeze/thawed 3x with liquid nitrogen with a 30 sec vortexing step between each freeze thaw cycle. After the final thaw, chloroform was added to a 1:5 chloroform/Trizol ratio and aqueous separation was performed using centrifugation in a heavy gel phase-lock tube (VWR, 10847-802). The aqueous phase was mixed 1:1 with isopropanol and then RNA was extracted as per manufacturer’s directions using a Qiagen RNeasy Mini Kit (74106). Library preparation was performed using a Kapa Biosystems mRNA Hyper Prep Kit as per manufacturer’s protocol and sequencing was performed at the Vincent J Coates Genomic Sequencing Core at the University of California, Berkeley using an Illumina HS4000 mode SR100. Three biological replicates were used per condition. Low quality reads and adaptor sequences were trimmed using Trim Galore v0.6.7-1(Krueger 2025). Reads were aligned to the WBcel235 genome using STAR-2.7.10a(Dobin et al. 2013a), and a raw count matrix was generated using featureCounts from Subread v2.0.3(Liao et al. 2019) and the WBcel235.107.gtf file. Differential gene expression was calculated with DESeq2(Love et al. 2014) and sva v3.42.0 was used to correct for batch effects and non-biological sources of variation(Leek et al. 2012). GO enrichment analysis was performed using enrichGO from clusterProfiler_4.14.6(Yu 2024).

## Results and Discussion

Previous studies have shown that heat-shock induces the UPR^ER^ in mammalian cells (Heldens et al. 2011). Here, we sought to determine whether heat-shock can also induce UPR^ER^ in vivo by testing the impact of heat-shock on the canonical UPR^ER^ reporter, *hsp-4p::GFP* in *C. elegans*, which is an indicator of *hsp-4*/HSPA5/BiP upregulation (Calfon et al. 2002). Indeed, the *hsp-4* promoter has predicted heat-shock responsive elements (Zha et al. 2019), suggesting potential activation of *hsp-4p::GFP* via HSF-1. We found significant activation of the *hsp-4p::GFP* reporter upon exposure to heat stress (**Fig. 1A-B**). Interestingly, this activation is dependent on both *hsf-1* and *xbp-1*, unlike UPR^ER^ induction via ER stress (exposure to tunicamycin, which blocks n-linked glycosylation and induces misfolded protein stress in the ER (Heifetz et al. 1979), which is dependent on *xbp-1*, but not on *hsf-1* (**Fig. S1**). Importantly, the induction of heat-induced UPR^ER^ is not dependent on the other branches of the UPR^ER^, as *pek-1* or *atf-6* RNAi had no effect (**Fig. 1C-D**).

**Fig. 1.**
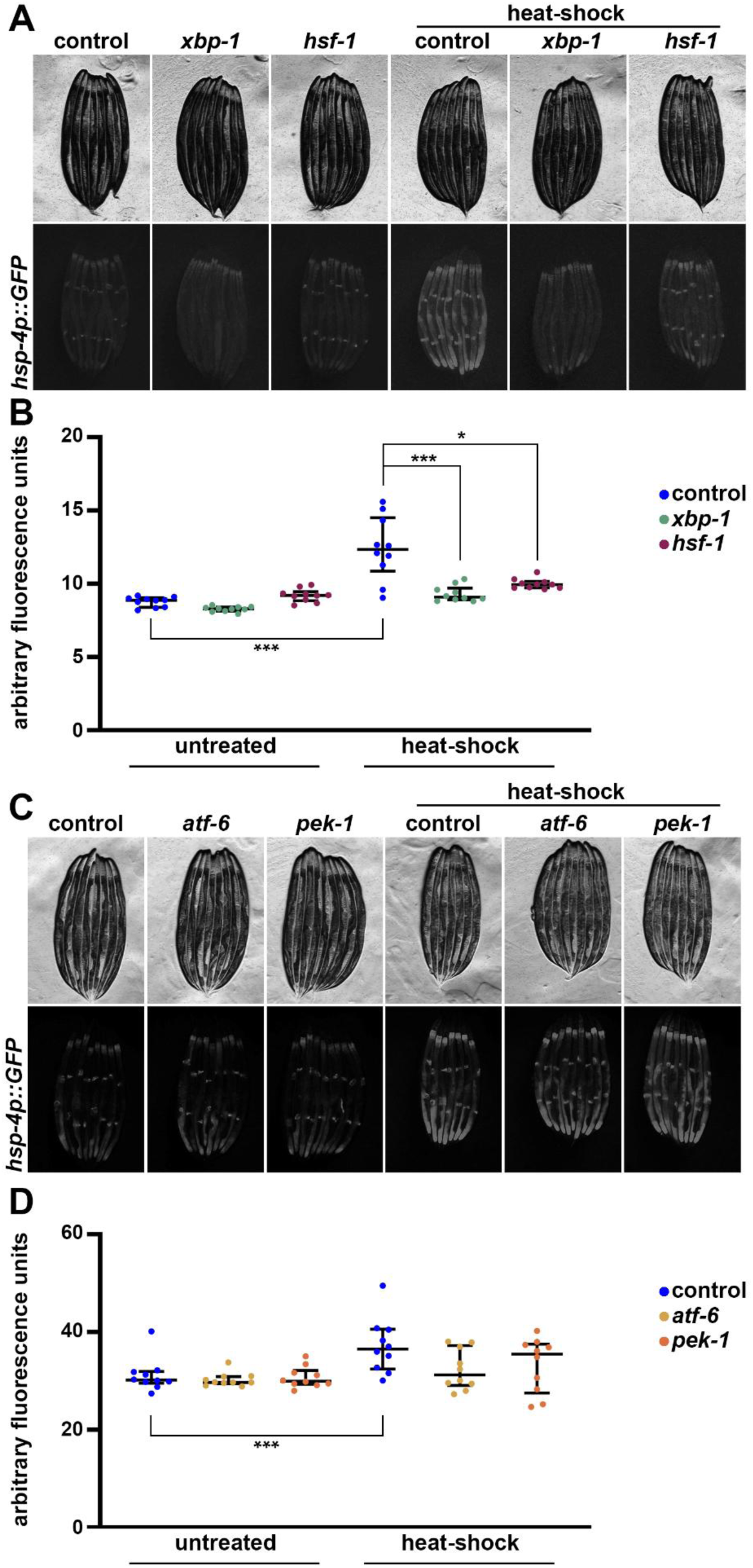
Heat stress induces UPR^ER^ in a *hsf-1* and *xbp-1* dependent manner. **(A)** Representative fluorescent images of day 1 adult animals expressing *hsp-4p::GFP* grown on empty vector (EV, control), *hsf-1*, or *xbp-1* RNAi bacteria from L1. Heat-shock conditions are 2 hours at 34 °C followed by a 4-hour recovery at 20 °C. Data is representative of 3 independent replicates. **(B)** Quantification of A represented as arbitrary fluorescent units, which are integrated fluorescence intensity measurements using Image J. Data is representative of three biological replicates where individual dots represent individual animals and lines represent median plus interquartile range. * = p < 0.05; *** = p < 0.001 using Kruskal-Wallis multiple comparison testing. **(C)** Representative fluorescent images similar to (A) but grown on EV (control), *xbp-1, atf-6,* or *pek-1* RNAi bacteria. **(D)** Quantification of (C) performed similar to (B).

To determine whether the activation of the UPR^ER^ under heat stress was distinct from that activated under ER stress conditions, we simultaneously exposed *hsp-4p::GFP* animals to both heat and ER stress. To apply ER stress, animals were grown on RNAi against *tag-335*, a putative mannose-1-phosphate guanylyltransferase whose knockdown results in robust UPR^ER^ activity (Higuchi-Sanabria, Shen, et al. 2020). Interestingly, we found that *tag-335* knockdown and heat-shock did not have additive effects on UPR^ER^ activation, suggesting that a maximal capacity of UPR^ER^ activation is achieved upon *tag-335* knockdown (**Fig. S2**). To further evaluate whether XBP-1 had any influence on heat-induced UPR^ER^, we next tested the impact of *xbp-1s* overexpression on UPR^ER^ activity. Consistent with previous reports, overexpression of *xbp-1s* in neuronal (Taylor and Dillin 2013; Daniele et al. 2020) or glial cells (Frakes et al. 2020) results in robust activation of UPR^ER^ (**Fig. 2**). Unlike UPR^ER^ activation via *tag-335* RNAi, UPR^ER^ induction via non-autonomous activation from glial or neuronal signals had additive effects with heat-induced UPR^ER^. These data suggest that either heat-induced or XBP-1-induced UPR^ER^ are distinct, or that *xbp-1s* overexpression can enhance heat-induced UPR^ER^ activity.

**Fig. 2.**
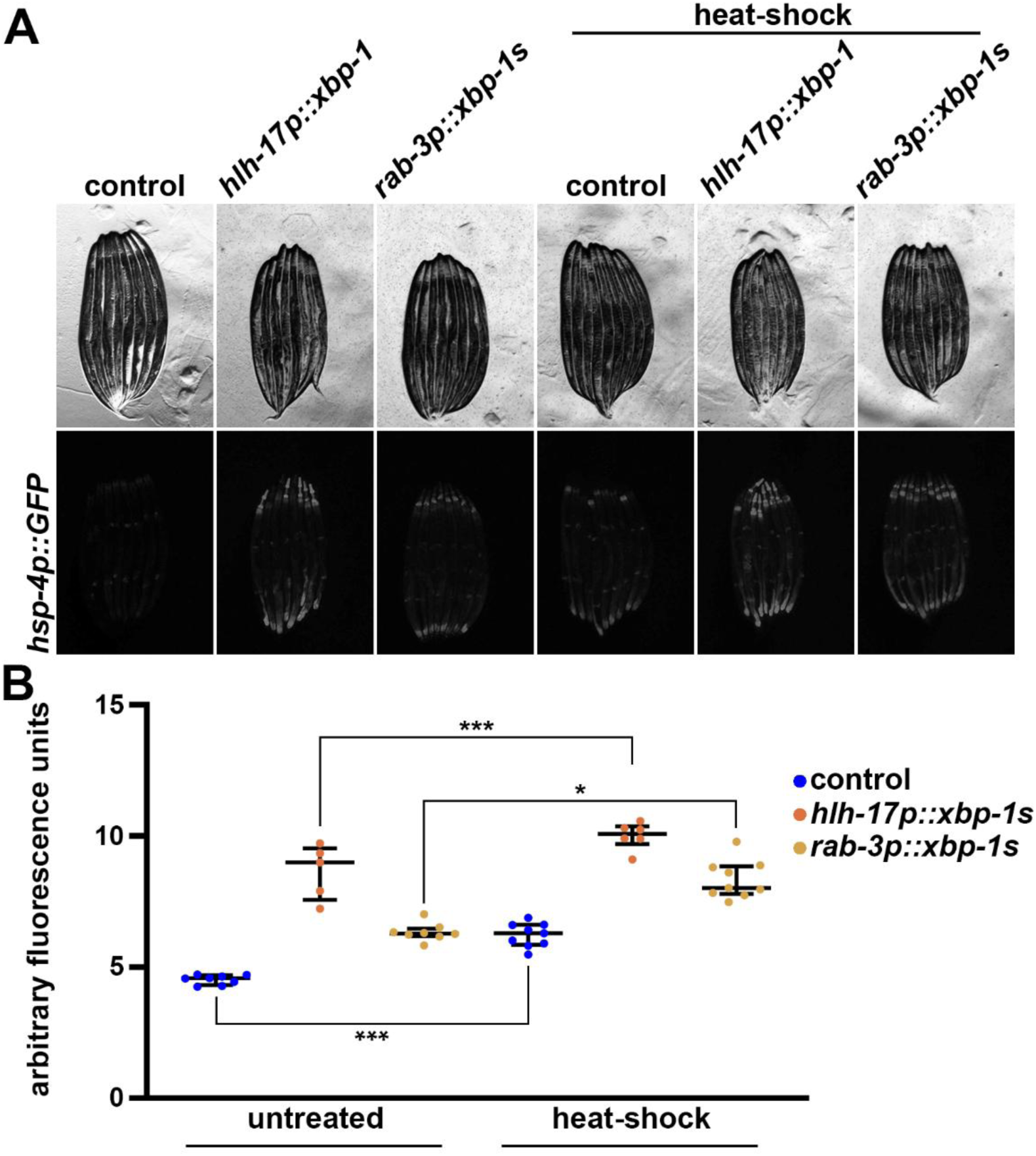
Neuronal or glial *xbp-1s* overexpression does not affect heat induced UPR^ER^. **(A)** Representative fluorescent images of day 1 adult animals expressing *hsp-4p::GFP* alone or in combination with *hlh-17p::xbp-1s* or *rab-3p::xbp-1s* grown on EV (control) bacteria from L1. Heat-shock conditions are 2 hours at 34 °C followed by a 4-hour recovery at 20 °C. Animals are imaged on day 1 of adulthood. Data is representative of 3 independent replicates. **(B)** Quantification of A represented as arbitrary fluorescent units, which are integrated fluorescence intensity measurements using Image J. Data is representative of three biological replicates where individual dots represent individual animals and lines represent median plus interquartile range. * = p < 0.05; *** = p < 0.001 using Kruskal-Wallis multiple comparison testing.

We next sought to determine whether regulators of UPR^ER^ could impact activity of the HSR. To measure the HSR, we utilized the canonical HSR reporter, *hsp-16.2p::GFP*, which has very low basal signal and robustly increases GFP expression upon exposure to heat stress (Link et al. 1999). To our surprise, we found that overexpression of *xbp-1s* in neurons or glial cells robustly enhanced the activity of the HSR, although it had no impact on basal *hsp-16.2p::GFP* expression (**Fig. 3**). Importantly, the increase in HSR induction upon exposure to heat stress is comparable to overexpression of *hsf-1* in neurons. To determine whether this increase in HSR had any physiological impact, we next measured whether *xbp-1s* overexpression could improve thermotolerance, a canonical feature of *hsf-1* overexpression (Baird et al. 2014). Indeed, neuronal or glial *xbp-1s* overexpression displayed a significant increase in thermotolerance, again comparable to neuronal *hsf-1* overexpression, similar to its effect on the HSR (**Fig. 4**), suggesting this increased activation of the HSR is physiologically relevant.

**Fig. 3.**
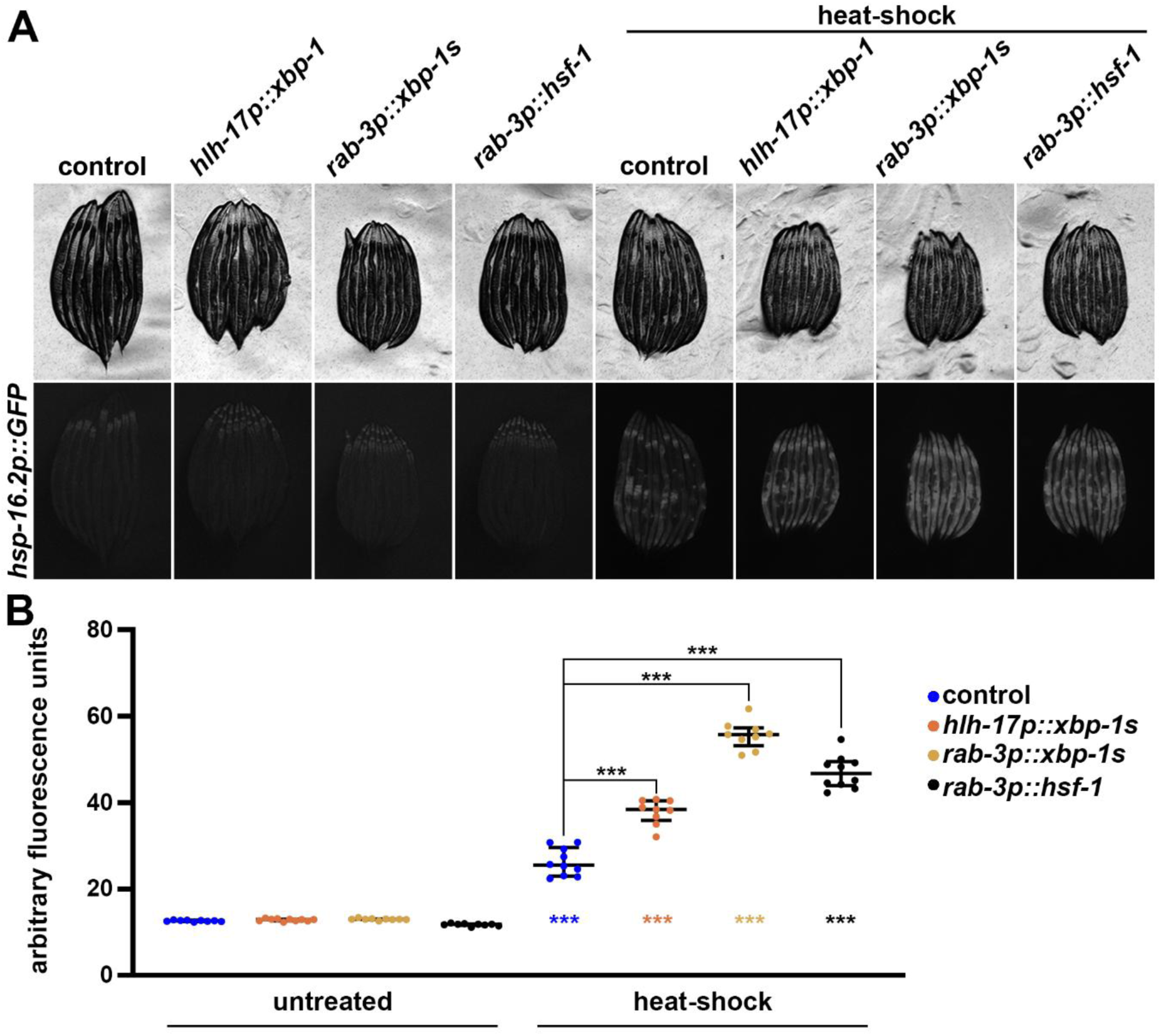
Neuronal and glial *xbp-1s* improves HSR similar to neuronal *hsf-1*. **(A)** Representative fluorescent images of day 1 adult animals expressing *hsp-16.2p::GFP* alone or in combination with *hlh-17p::xbp-1s, rab-3p::xbp-1s,* or *rab-3p::hsf-1* grown on EV (control) bacteria from L1. Heat-shock conditions are 2 hours at 34 °C followed by a 2-hour recovery at 20 °C. Animals are imaged on day 1 of adulthood. Data is representative of 3 independent replicates. **(B)** Quantification of A represented as arbitrary fluorescent units, which are integrated fluorescence intensity measurements using Image J. Data is representative of three biological replicates where individual dots represent individual animals and lines represent median plus interquartile range. *** = p < 0.001 using Kruskal-Wallis multiple comparison testing. Colored *** below each plot indicate p-value comparing heat-shocked animals vs. their untreated control counterpart.

**Fig. 4.**
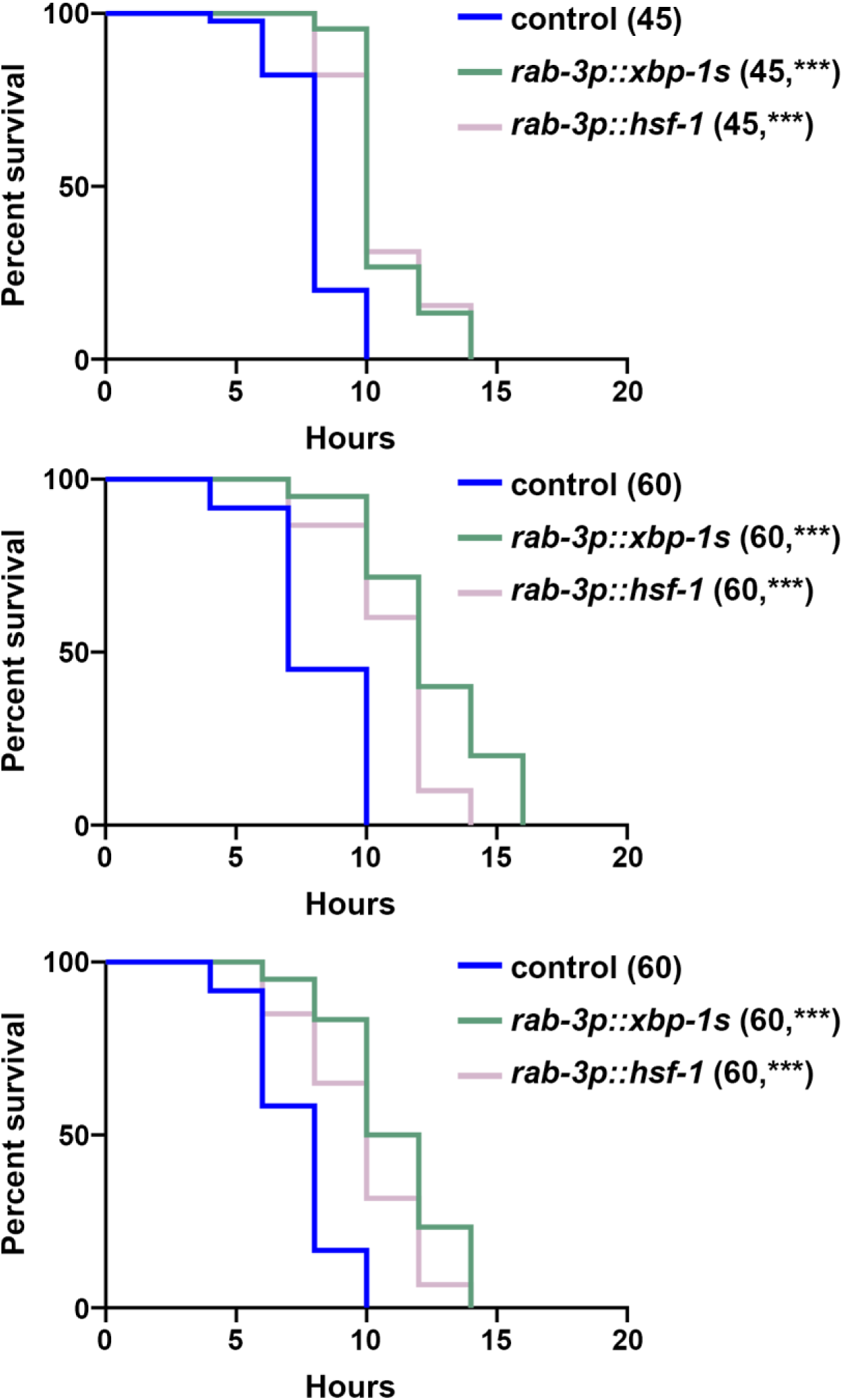
Neuronal *xbp-1s* improves thermotolerance similar to neuronal *hsf-1*. N2 wild-type (control), *rab-3p::xbp-1s*, or *rab-3p::hsf-1* animals grown on EV RNAi bacteria from L1. Animals were moved to 34 °C on day 1 of adulthood and survival was scored every 2 hours. All three biological replicates are shown for full transparency as thermotolerance measurements have been shown to display significant variability by several groups (Zevian and Yanowitz 2014; Bar-Ziv et al. 2020; Torres et al. 2022). Sample size is represented in the legend in parentheses and *** = p < 0.0001 vs. control using Log-Rank testing in PRISM 10.

Although the usage of transcriptional reporters allows for robust and rapid characterization of numerous conditions, it suffers from several caveats and limitations. First, GFP expression does not directly correlate with endogenous gene expression, as GFP stability is not directly correlated with transcript or protein stability of the actual gene of interest. In addition, all the transcriptional reporters presented here are overexpression constructs integrated in random gene loci, and thus the regulation of gene expression is potentially not identical to the endogenous gene construct. Finally, the overreliance on a single gene reporter to define a complex stress response that involves dozens to hundreds of transcriptional changes can be inherently flawed. Indeed, a recent study (Kim et al. 2025) has shown that the overreliance on the *hsp-6p::GFP* UPR^MT^ reporter may have been a contributing cause to the controversy linking UPR^MT^ activity to longevity in *C. elegans* (Bennett et al. 2014). Therefore, to more globally understand the heat-induced UPR^ER^, we performed transcriptomic analysis.

Here, we performed RNA sequencing on wild-type and neuronal *hsf-1* animals. For neuronal *hsf-1*, we compared overexpression of full-length *hsf-1* (labeled FL for the transcriptomics section to avoid confusion) and a c-terminal deletion (CTD) of *hsf-1*, which has been shown to maintain its ability to activate targets of the actin cytoskeleton, while losing its capacity to induce the HSR (Baird et al. 2014). We also performed RNA-sequencing on the *hsf-1(sy441)* mutant, which carries the same CTD, which allows interrogation of HSF-1 activity in the absence of HSR activation. We performed bulk RNA sequencing on >800 animals per condition using three biological replicates per condition. After library construction, we performed FastQC analysis on each library using MultiQC (Ewels et al. 2016). All libraries displayed similar Phred scores >35, suggesting there was no systemic bias favoring any single condition and high quality of our libraires (**Fig. S3A**). Sequencing results were mapped to the *C. elegans* reference genome WBcel235 using the STAR tool, which mapped sufficient and consistent unique reads across all samples (**Fig. S3B**) (Dobin et al. 2013b; Howe et al. 2021). Each biological replicate shared similar expression profiles as indicated by similar Spearman correlation (**Fig. S3C**) and close clustering using principal component analysis (**Fig. S3D**). As expected, the untreated groups clustered closely together, as did the heat-shocked conditions. Interestingly, the *hsf-1(sy441)* clustered the furthest away from all other samples, suggesting the mutation of *hsf-1* has the most robust impact on bulk transcriptome both under basal conditions and heat-shock.

Consistent with our *hsp-4p::GFP* reporter data, wild-type animals treated with heat-shock resulted in a significant upregulation of *hsp-4* transcripts (**Fig. 5A**). Heat-shock also increased transcript levels of *xbp-1*, which provides further evidence that XBP-1-mediated UPR^ER^ is activated under conditions of heat-shock. Indeed, further analysis revealed that at least a handful of XBP-1 targets are upregulated under heat-shocked conditions (**Fig. 5B**). Interestingly, *hsf-1(sy441)* animals still displayed a similar increase in *hsp-4* and *xbp-1* transcripts, as well as increased XBP-1 targets, suggesting that the canonical HSR mediated by a full-length HSF-1 is not required for heat-induced UPR^ER^. Importantly, gene ontology (GO) analysis of all significantly upregulated genes of heat-shocked vs. untreated animals for each group showed an expected increase in genes associated with protein folding, chaperone function, and other HSR targets, which was lost in the *hsf-1(sy441)*, showing further evidence that our transcriptomic results were robust and reproduced previous findings (Baird et al. 2014) (**Fig. 6**). Interestingly, the *hsf-1FL* included the UPR^ER^ as one of the top 15 enriched GO terms, although this GO term was significant in all conditions (**Table S1**). Our transcriptome analysis further substantiates the existence of a heat-induced UPR^ER^ that is likely dependent on both *xbp-1* and *hsf-1*.

**Fig. 5.**
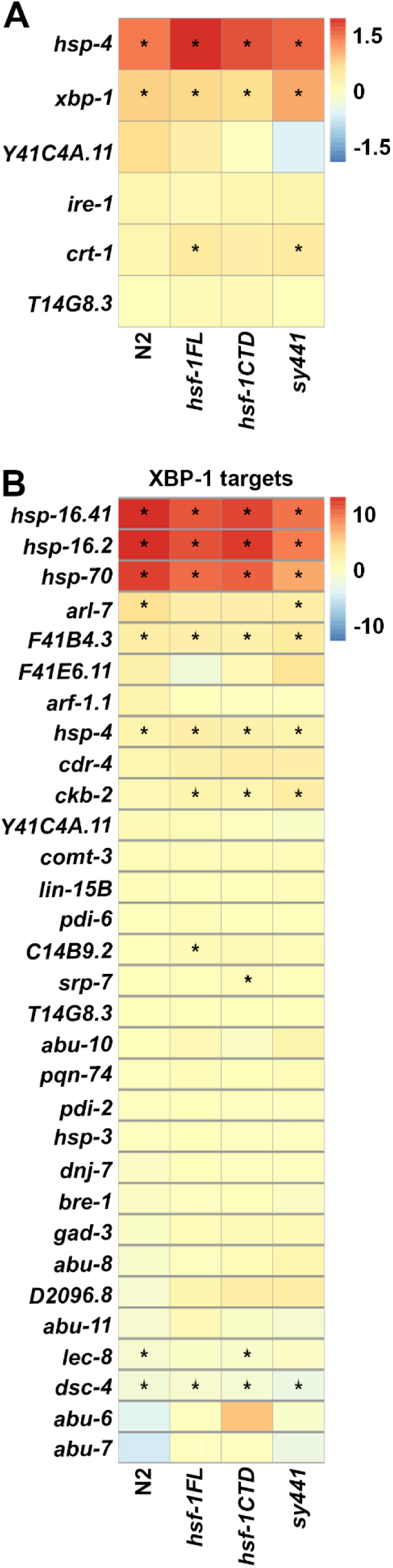
Heat-shock induces UPR^ER^ genes. RNA sequencing was performed in N2, *rab-3p::hsf-1FL*, *rab-3p::hsf-1CTD*, and *hsf-1(sy441)*, animals with and without heat-shock. DEGs were identified by comparing each heat-shock condition against untreated and are available in **Table S2**. **(A)** Heat-map of log_2_(fold-change) of N2, *rab-3p::hsf-1FL*, *rab-3p::hsf-1CTD*, and *hsf-1(sy441)* for canonical UPR^ER^ genes, *hsp-4*, *xbp-1*, *Y41C4A.11*, *ire-1*, *crt-1*, and *T14G8.3*. Warmer colors indicate increased expression and cooler colors indicate decreased expression. * = p < 0.01. See **Table S3** for gene list. **(B)** Heat-map of log_2_(fold-change) of XBP-1 targets.

**Fig. 6.**
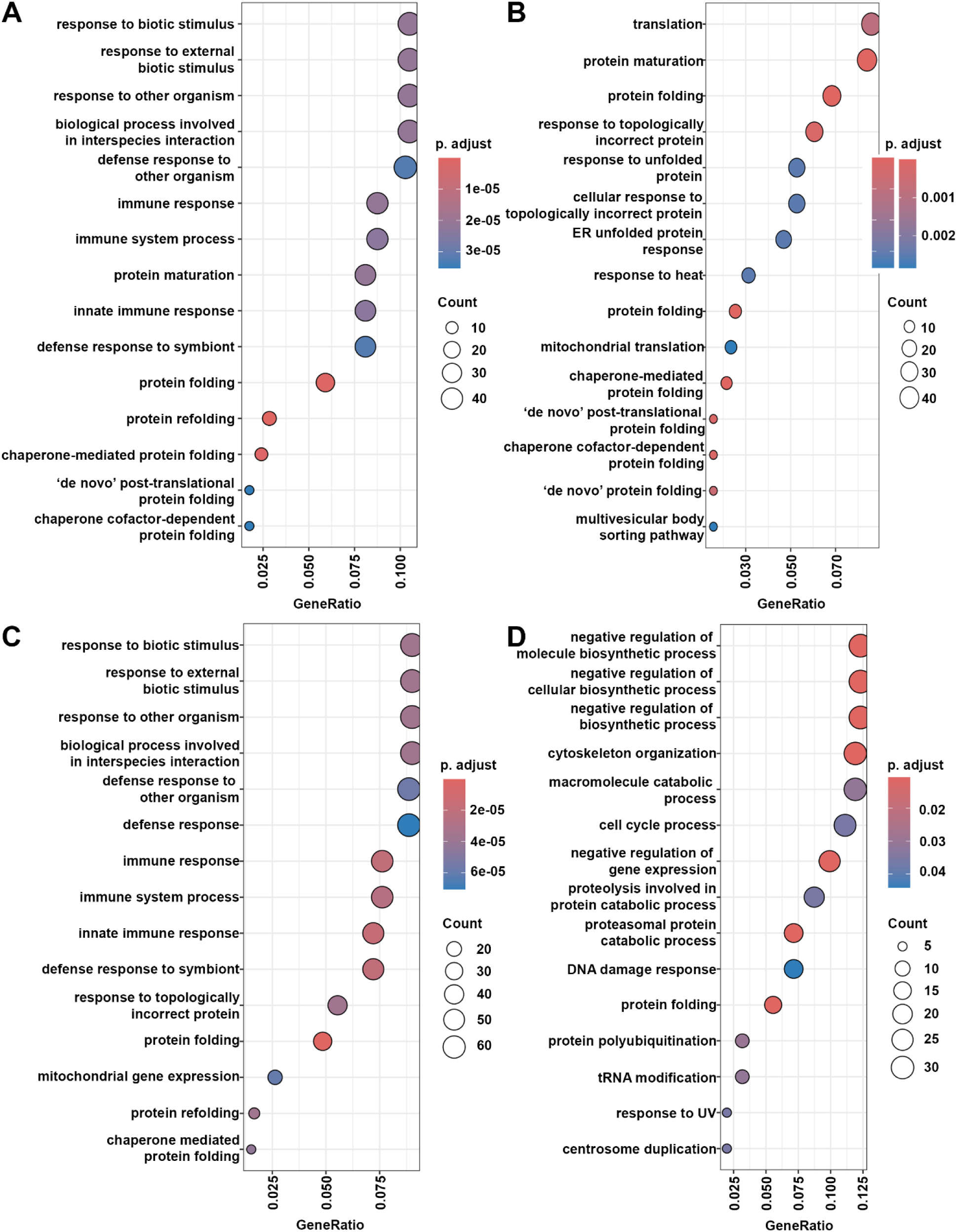
GO analysis of heat-shock induced genes. **(A)** Top 15 GO-terms for N2 heat-shock vs. N2 untreated. **Table S4** for gene list. **(B)** Top 15 GO-terms for *rab-3p::hsf-1FL* heat-shock vs. *rab-3p::hsf-1FL* untreated. **(C)** Top 15 GO-terms for *rab-3p::hsf-1CTD* heat-shock vs. *rab-3p::hsf-1CTD* untreated. **(D)** Top 15 GO terms for *hsf-1(sy441)* heat-shock vs. *hsf-1(sy441)* untreated. A table of all significant GO terms is available in **Table S1**. All DEGs of each condition against N2 controls are also available in **Table S5**.

## Discussion

Overall, our findings highlight the complex interplay between the HSR and the UPR^ER^ in *C. elegans*, emphasizing the existence of both compartment-specific and shared regulatory mechanisms. The observation that there exists a heat-induced HSF-1 and XBP-1 dependent UPR^ER^ distinct from the canonical UPR^ER^ mediated primarily by XBP-1 underscores the plasticity of stress response pathways. This adds to the growing literature of cross communication of stress responses, including prior work showing that cytosolic HSF-1 can influence mitochondrial proteostasis through fatty acid remodeling (Kim et al. 2016) and that mitochondrial stress can delay the age-associated loss of HSR activity (Labbadia et al. 2017). Similarly, our data suggest that cytosolic HSF-1 may directly – or indirectly – modulate ER stress response, though the exact mechanism of transcriptional activation and shared chaperone networks remains to be elucidated. One interesting future avenue of research would be to understand the connection between HSF-1 and XBP-1. We found that heat-induced UPR^ER^ requires both transcription factors, but whether heat stress induces direct interaction of HSF-1 and XBP-1 or an indirect, coordinated effort to regulate transcription remains to be explored. Indeed, XBP-1 has been shown to interact with other stress response factors (Glover-Cutter et al. 2013), such as SKN-1 to regulate both ER and oxidative stress responses, suggesting interaction with other stress response transcription factors is possible.

The bidirectional crosstalk between the UPR^ER^ and HSR adds complexity to the rapidly growing understanding of stress response integration. The integrated stress response (ISR) serves as a well-characterized framework for understanding the integration of multiple stress inputs into a shared response. The ISR responds to a diverse set of stressors including proteotoxic stress, nutrient deprivation, organelle stress (Kalinin et al. 2023; Wek et al. 2023; Lu et al. 2024; Ryoo 2024). The ISR typically involves eIF2α phosphorylation via kinases like PEK-1 to reduce global translation and reduce translational burden under stress. Parallel to the reduction of translation via the ISR, our study highlights a convergent response to variable stressors to promote proteostasis. Our study finds that heat stress results in induction of chaperones required for both cytosolic and ER function, akin to how mitochondrial stress preconditions the HSR (Labbadia et al. 2017), inhibition of cytosolic chaperones promotes activation of the UPR^MT^ (Kim et al. 2016), and how loss of cytosolic proteostasis can be offset by mitochondrial proteostasis (Berendzen et al. 2016; Wang et al. 2024). These phenomena may reflect a general strategy wherein localized stress can activate numerous proteostasis pathways to prevent cellular damage.

This converging stress responsive pathway is further strengthened through inter-tissue communication of stress. Our study adds to the growing understanding of non-autonomous signaling of stress by highlighting the ability of neuronal and glial *xbp-1s* overexpression to enhance HSR activity and thermotolerance, despite XBP-1 being dispensable for the HSR. This points to a heterotypic neuronal signaling mechanism. This mirrors earlier findings whereby neuronal *hsf-1* activates both HSF-1 and DAF-16 in peripheral tissues to activate both HSR and OxSR to extend longevity (Douglas et al. 2015). Whether neural *xbp-1s* similarly regulates a heterotypic response has not yet been explored, but neuronal *xbp-1s* has been shown to activate several distinct peripheral responses, including improved proteostasis (Taylor and Dillin 2013), lipid homeostasis (Imanikia, Sheng, et al. 2019; Daniele et al. 2020), autophagy (Imanikia, Özbey, et al. 2019), and immune response (Coakley et al. 2025). Similarly, glial *xbp-1s* also mediates a peripheral proteostasis (Frakes et al. 2020), lipid metabolism, and autophagy response (Metcalf et al. 2024). Importantly, both neuronal and glial *xbp-1s* lifespan extension required both peripheral XBP-1 and HLH-30 (Imanikia, Özbey, et al. 2019; Metcalf et al. 2024), suggesting a similar heterotypic response. Our data suggests that neural *xbp-1s* also amplifies the HSR in the periphery to promote thermotolerance. Still to be determined is whether *hsf-1* is required for the other beneficial effects of neural *xbp-1s*, considering its previously established roles in proteostasis (Morley and Morimoto 2004; Hayashida et al. 2010), autophagy (Kumsta et al. 2017), lipid metabolism (Chauve et al. 2021; Watterson et al. 2022), and longevity (Hsu et al. 2003; Morley and Morimoto 2004).

In mammals, hypothalamic *Xbp1s* activation improved hepatic metabolism, suggesting conserved principles of inter tissue coordination (Williams et al. 2014). Whether these non-autonomous signals are also dependent on a heterotypic response including activation of HSF-1 or other transcriptional regulators is uncertain. However, human cancer cells activate an atypical UPR^ER^ activation upon exposure to heat stress, which was independent of canonical ER stress pathways like PERK (Heldens et al. 2011). Importantly, activated targets included ER chaperones such as HSPA5/BiP, the target of our *hsp-4p::GFP* reporter, suggesting relevance of our findings to a mammalian context. Overall, our work advances the understanding of the interconnectivity of stress responses and further substantiates *C. elegans* as a powerful in vivo model system for dissecting conserved mechanisms of systemic stress coordination. Future work is critical to focus on identifying the signaling mechanisms of the heterotypic responses of neural UPR to peripheral HSF-1 and understanding the potential interaction between XBP-1 and HSF-1 to coordinate both ER and heat stress responses to converge on shared downstream effectors to promote resilience and longevity.

## Data Availability Statement

All data are presented in the figures and tables of the manuscript. All RNA-sequencing data reanalyzed from previously published datasets and raw data are available at Annotare E-MTAB-9771 (Higuchi-Sanabria, Durieux, et al. 2020) and the National Center for Biotechnology Information Sequence Read Archive repository accession number BioProject PRJNA589459 (Frakes et al. 2020). RNA-sequencing data produced here are available at National Center for Biotechnology Information Sequence Read Archive repository accession number PRJNA1399626.

## Acknowledgements

We thank the Douglas lab for sharing RNA-seq datasets. We thank the Dillin lab for sharing reagents and resources.

## Author Contributions

T.C.T., A.A., G.G., and R.H.S. collaboratively contributed to the study design, experiments, data analysis, preparation of figures, and writing the manuscript. R.A.B. assisted with quantification of imaging data. P.A.F. assisted with RNA-seq analysis.

---

**Fig. S1.**
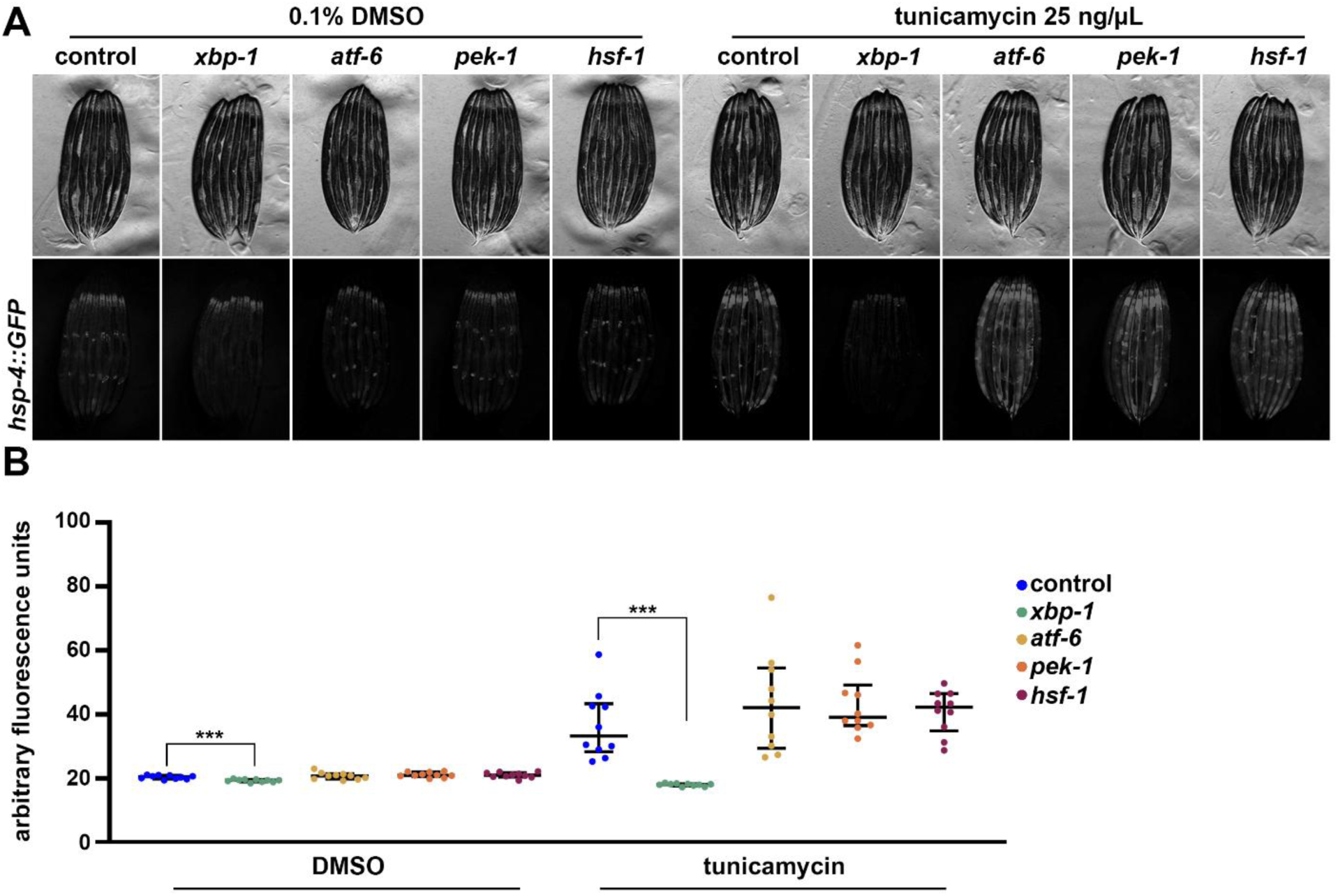
*hsf-1* is not required for ER stress induced UPR^ER^. **(A)** Representative fluorescent images of day 1 adult animals expressing *hsp-4p::GFP* grown on EV (control), *xbp-1, atf-6, pek-1,* or *hsf-1* RNAi bacteria from L1. L4 animals are treated with 0.1% DMSO or 25 ng/µL tunicamycin in M9 solution in a rotator at 20 °C for 4 hours. DMSO or tunicamycin are washed 3x with M9 solution and animals are placed on standard RNAi plates with EV RNAi bacteria and grown at 20 °C to recover overnight (maximum of 16 hours). Animals are imaged on day 1 of adulthood. Data is representative of 3 independent replicates. **(B)** Quantification of A represented as arbitrary fluorescent units, which are integrated fluorescence intensity measurements using Image J. Data is representative of three biological replicates where individual dots represent individual animals and lines represent median plus interquartile range. *** = p < 0.001 using Kruskal-Wallis multiple comparison testing.

**Fig. S2.**
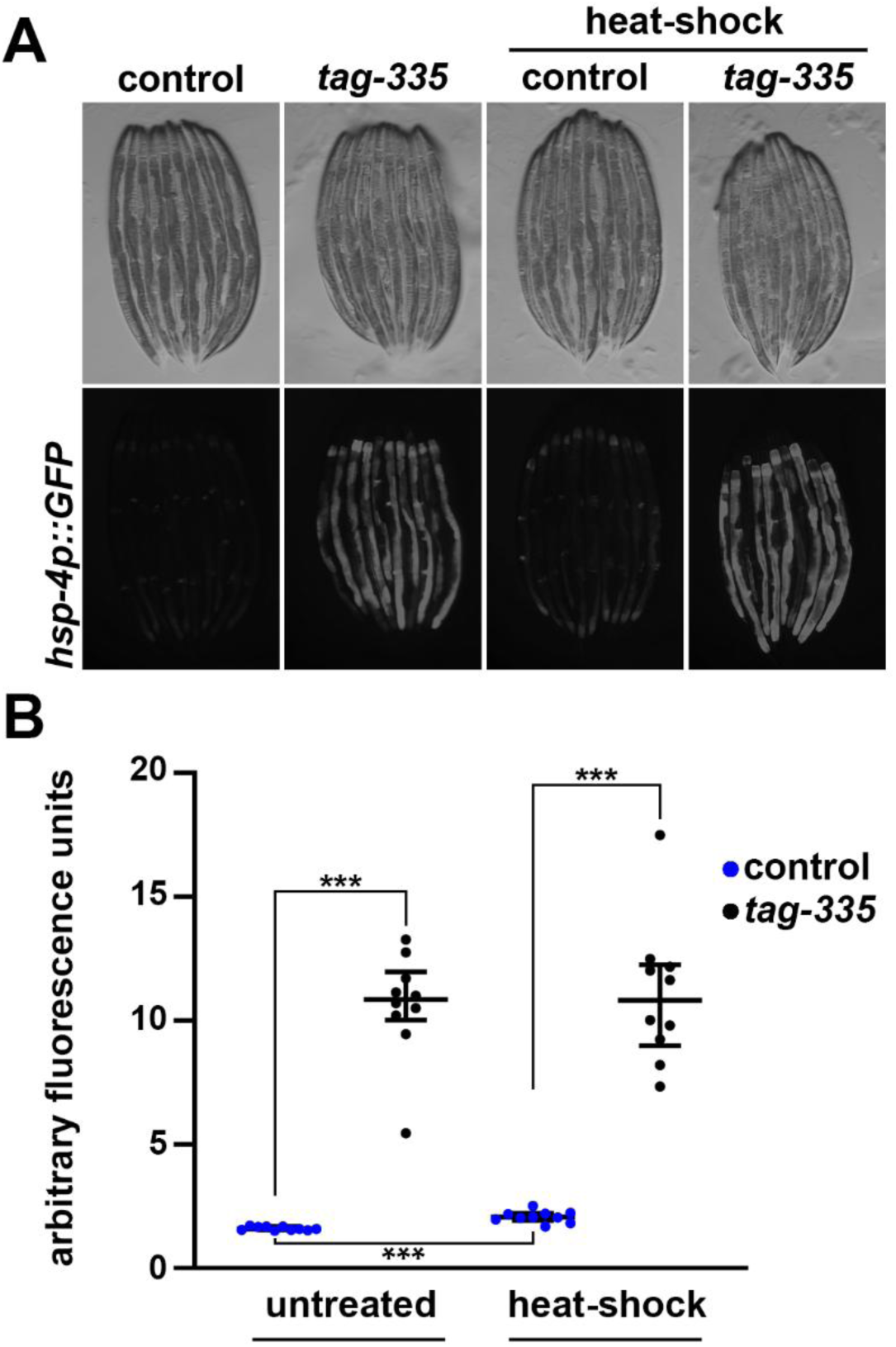
ER and heat stress mediated UPR^ER^ are not additive. **(A)** Representative fluorescent images of day 1 adult animals expressing *hsp-4p::GFP* grown on EV (control) or *tag-335* RNAi bacteria from L1. Heat-shock conditions are 2 hours at 34 °C followed by a 4-hour recovery at 20 °C. Animals are imaged on day 1 of adulthood. Data is representative of 3 independent replicates. **(B)** Quantification of A represented as arbitrary fluorescent units, which are integrated fluorescence intensity measurements using Image J. Data is representative of three biological replicates where individual dots represent individual animals and lines represent median plus interquartile range. *** = p < 0.001 using Kruskal-Wallis multiple comparison testing.

**Fig. S3.**
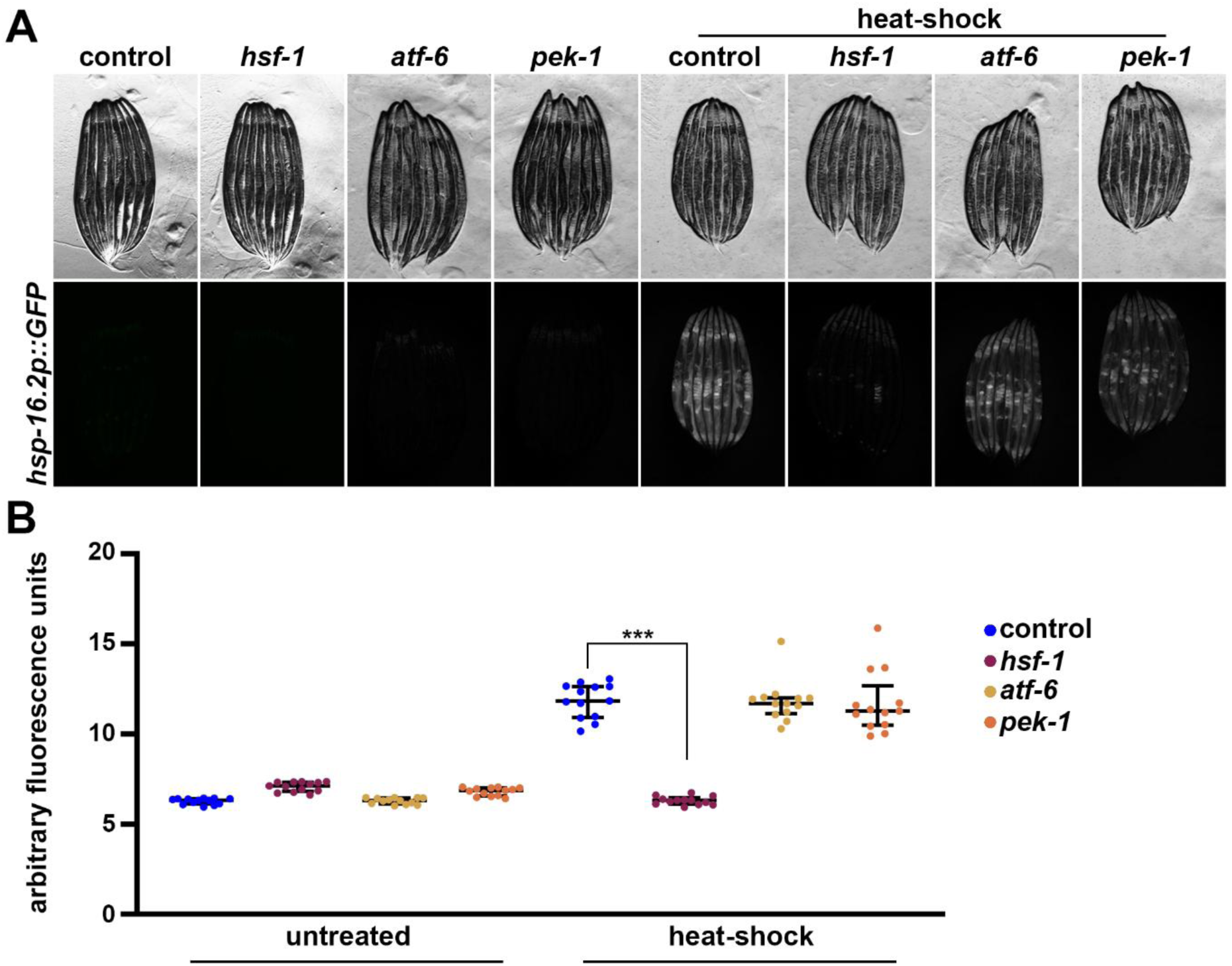
The HSR requires *pek-1* and *atf-6.* **(A)** Representative fluorescent images of day 1 adult animals expressing *hsp-16.2p::GFP* grown on EV (control), *hsf-1, atf-6,* or *pek-1* RNAi bacteria from L1. Heat-shock conditions are 2 hours at 34 °C followed by a 2-hour recovery at 20 °C. Animals are imaged on day 1 of adulthood. Data is representative of 3 independent replicates. **(B)** Quantification of A represented as arbitrary fluorescent units, which are integrated fluorescence intensity measurements using Image J. Data is representative of three biological replicates where individual dots represent individual animals and lines represent median plus interquartile range. * = p < 0.05; *** = p < 0.001 using Kruskal-Wallis multiple comparison testing.

**Fig. S4.**
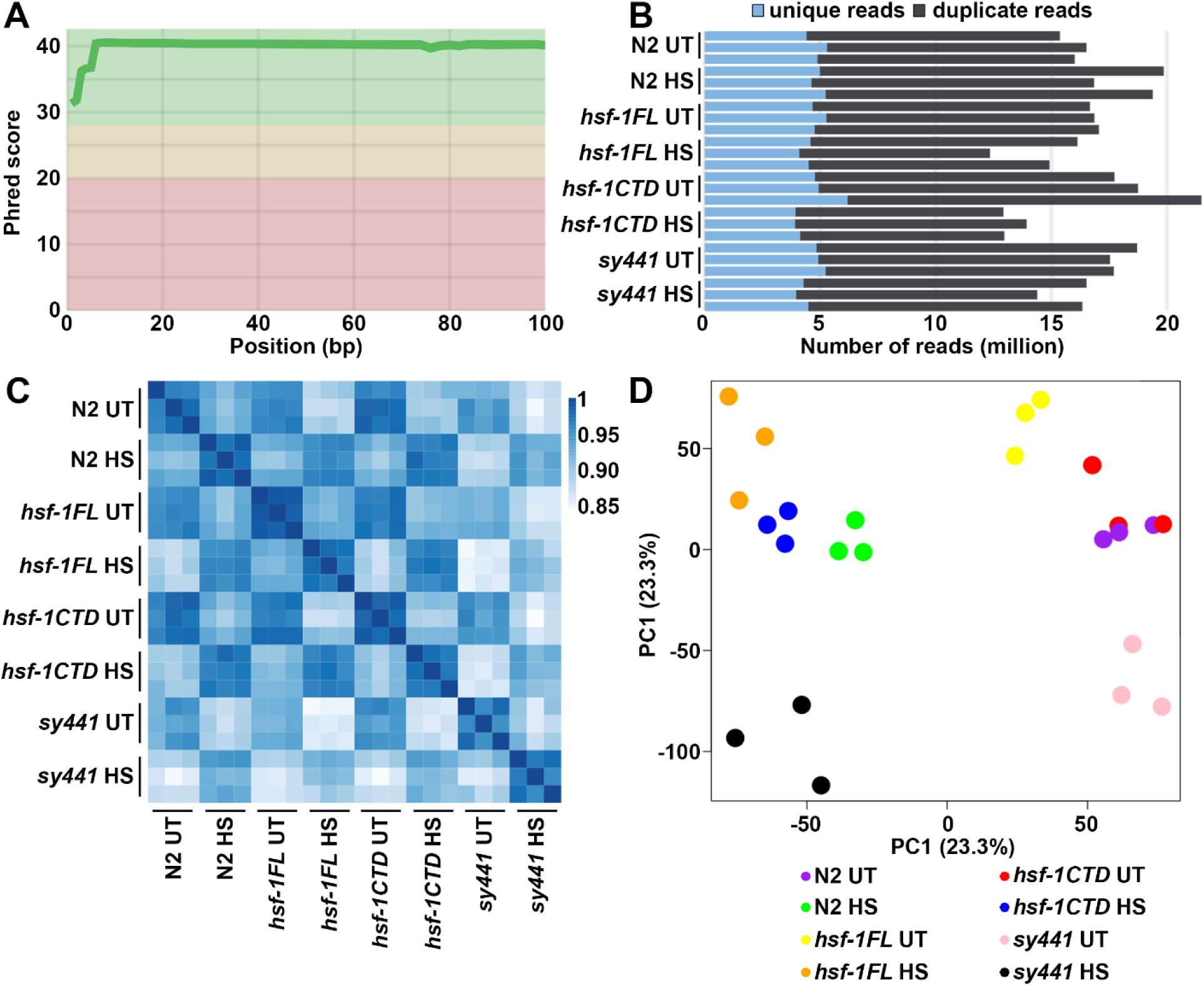
Quality control for RNA-seq libraries of heat-shock activation in *C. elegans*. **(A)** Mean quality score (Phred score) of each RNA sequencing library. X-axis indicates the base pair (bp) position of each sequence and the y-axis indicates the Phred score. The graph was generated by MultiQC(Ewels et al. 2016). **(B)** The number of unique (blue) and duplicated (black) reads from each pair-wise sequencing library (n = 3). **(C)** Spearman correlation plot of all RNA sequencing libraries (n = 3). **(D)** PCA plots of all RNA sequencing libraries.

**Table S1. Table of all significant GO terms.** All significant GO terms for for N2, *rab-3p::hsf-1FL*, *rab-3p::hsf-1CTD*, and *hsf-1(sy441)* animals for all significant DEGs of heat-shocked vs. untreated controls for each condition.

**Table S2. DEGs of HS vs. UT.** All differentially expressed genes represented as log_2_(fold-change) for N2, *rab-3p::hsf-1FL*, *rab-3p::hsf-1CTD*, and *hsf-1(sy441)* animals calculated as heat-shocked vs. untreated controls for each condition.

**Table S3 DEGs for UPR heat map.** All log_2_(fold-change) used to create heat-map of **Fig. 5A**.

**Table S4. DEGs for XBP-1 targets heat map.** All log_2_(fold-change) used to create heat-map of **Fig. 5B**.

**Table S5. DEGs against N2 control.** All differentially expressed genes represented as log_2_(fold-change) N2, *rab-3p::hsf-1FL*, *rab-3p::hsf-1CTD*, and *hsf-1(sy441)* animals with heat-shock (HS) or left untreated (UT) calculated against N2 controls.

## References

Anckar J, Sistonen L. 2011. Regulation of HSF1 Function in the Heat Stress Response: Implications in Aging and Disease. 80(1):1089–1115. 10.1146/annurev-biochem-060809-095203

Baird NA et al. 2014. HSF-1-mediated cytoskeletal integrity determines thermotolerance and life span. Science. 346(6207):360–363. 10.1126/science.1253168

Bar-Ziv R et al. 2020. Measurements of Physiological Stress Responses in C. Elegans. JoVE (Journal of Visualized Experiments). (159):e61001. 10.3791/61001

B’chir W et al. 2013. The eIF2α/ATF4 pathway is essential for stress-induced autophagy gene expression. Nucleic Acids Res. 41(16):7683–7699. 10.1093/nar/gkt563

Bennett CF et al. 2014. Activation of the mitochondrial unfolded protein response does not predict longevity in Caenorhabditis elegans. Nat Commun. 5(1):3483. 10.1038/ncomms4483

Berendzen KM et al. 2016. Neuroendocrine Coordination of Mitochondrial Stress Signaling and Proteostasis. Cell. 166(6):1553–1563.e10. 10.1016/j.cell.2016.08.042

Caldwell SR, Hill KJ, Cooper AA. 2001. Degradation of endoplasmic reticulum (ER) quality control substrates requires transport between the ER and Golgi. J Biol Chem. 276(26):23296–23303. 10.1074/jbc.M102962200

Calfon M et al. 2002. IRE1 couples endoplasmic reticulum load to secretory capacity by processing the XBP-1 mRNA. Nature. 415(6867):92–96. 10.1038/415092a

Chauve L et al. 2021. Neuronal HSF-1 coordinates the propagation of fat desaturation across tissues to enable adaptation to high temperatures in C. elegans. PLOS Biology. 19(11):e3001431. 10.1371/journal.pbio.3001431

Chen EY et al. 2013. Enrichr: interactive and collaborative HTML5 gene list enrichment analysis tool. BMC Bioinformatics. 14(1):128. 10.1186/1471-2105-14-128

Coakley AJ et al. 2025. Distinct responses to non-autonomous UPRER mediated by glutamatergic and octopaminergic neurons. Commun Biol. 8(1):1650. 10.1038/s42003-025-09036-1

Dai C. 2018. The heat-shock, or HSF1-mediated proteotoxic stress, response in cancer: from proteomic stability to oncogenesis. Philosophical transactions of the Royal Society of London Series B, Biological sciences. 373(1738). 10.1098/rstb.2016.0525

Daniele JR et al. 2020. UPRER promotes lipophagy independent of chaperones to extend life span. Science Advances. 6(1):eaaz1441. 10.1126/sciadv.aaz1441

Dobin A et al. 2013a. STAR: ultrafast universal RNA-seq aligner. Bioinformatics. 29(1):15–21. 10.1093/bioinformatics/bts635

Dobin A et al. 2013b. STAR: ultrafast universal RNA-seq aligner. Bioinformatics. 29(1):15–21. 10.1093/bioinformatics/bts635

Douglas PM et al. 2015. Heterotypic Signals from Neural HSF-1 Separate Thermotolerance from Longevity. Cell Rep. 12(7):1196–1204. 10.1016/j.celrep.2015.07.026

Ewels P, Magnusson M, Lundin S, Käller M. 2016. MultiQC: summarize analysis results for multiple tools and samples in a single report. Bioinformatics. 32(19):3047–3048. 10.1093/bioinformatics/btw354

Frakes AE et al. 2020. Four glial cells regulate ER stress resistance and longevity via neuropeptide signaling in C. elegans. Science. 367(6476):436–440. 10.1126/science.aaz6896

Frakes AE, Dillin A. 2017. The UPR(ER): Sensor and Coordinator of Organismal Homeostasis. Mol Cell. 66(6):761–771. 10.1016/j.molcel.2017.05.031

Glover-Cutter KM, Lin S, Blackwell TK. 2013. Integration of the Unfolded Protein and Oxidative Stress Responses through SKN-1/Nrf. PLOS Genetics. 9(9):e1003701. 10.1371/journal.pgen.1003701

Han J et al. 2013. ER-stress-induced transcriptional regulation increases protein synthesis leading to cell death. Nat Cell Biol. 15(5):481–490. 10.1038/ncb2738

Hayashida N et al. 2010. Heat shock factor 1 ameliorates proteotoxicity in cooperation with the transcription factor NFAT. The EMBO Journal. 29(20):3459–3469. 10.1038/emboj.2010.225

Haze K et al. 1999. Mammalian transcription factor ATF6 is synthesized as a transmembrane protein and activated by proteolysis in response to endoplasmic reticulum stress. Mol Biol Cell. 10(11):3787–3799. 10.1091/mbc.10.11.3787

Heifetz A, Keenan RW, Elbein AD. 1979. Mechanism of action of tunicamycin on the UDP-GlcNAc:dolichyl-phosphate Glc-NAc-1-phosphate transferase. Biochemistry. 18(11):2186–2192. 10.1021/bi00578a008

Heldens L et al. 2011. An atypical unfolded protein response in heat shocked cells. PLoS ONE. 6(8):e23512. 10.1371/journal.pone.0023512

Higuchi-Sanabria R, Shen K, et al. 2020. Lysosomal recycling of amino acids affects ER quality control. Science Advances. 6(26):eaaz9805. 10.1126/sciadv.aaz9805

Higuchi-Sanabria R, Durieux J, et al. 2020. Divergent Nodes of Non-autonomous UPRER Signaling through Serotonergic and Dopaminergic Neurons. Cell Rep. 33(10):108489. 10.1016/j.celrep.2020.108489

Howe KL et al. 2021. Ensembl 2021. Nucleic Acids Research. 49(D1):D884–D891. 10.1093/nar/gkaa942

Hsu A-L, Murphy CT, Kenyon C. 2003. Regulation of aging and age-related disease by DAF-16 and heat-shock factor. Science. 300(5622):1142–1145. 10.1126/science.1083701

Imanikia S, Sheng M, et al. 2019. XBP-1 Remodels Lipid Metabolism to Extend Longevity. Cell Reports. 28(3):581–589.e4. 10.1016/j.celrep.2019.06.057

Imanikia S, Özbey NP, et al. 2019. Neuronal XBP-1 Activates Intestinal Lysosomes to Improve Proteostasis in C. elegans. Curr Biol. 29(14):2322–2338.e7. 10.1016/j.cub.2019.06.031

Kalinin A, Zubkova E, Menshikov M. 2023. Integrated Stress Response (ISR) Pathway: Unraveling Its Role in Cellular Senescence. Int J Mol Sci. 24(24):17423. 10.3390/ijms242417423

Kim H-E et al. 2016. Lipid Biosynthesis Coordinates a Mitochondrial-to-Cytosolic Stress Response. Cell. 166(6):1539–1552.e16. 10.1016/j.cell.2016.08.027

Kim J, Dutta N, Garcia G, Higuchi-Sanabria R. 2025. Transcriptomic analysis of mitohormesis associated with lifespan extension in Caenorhabditis elegans. 2025.04.15.648933 [accessed 2025 Apr 25]. https://www.biorxiv.org/content/10.1101/2025.04.15.648933v1. 10.1101/2025.04.15.648933

Kohno K et al. 1993. The promoter region of the yeast KAR2 (BiP) gene contains a regulatory domain that responds to the presence of unfolded proteins in the endoplasmic reticulum. Mol Cell Biol. 13(2):877–890

Krueger F. 2025. FelixKrueger/TrimGalore. [accessed 2025 Apr 13]. https://github.com/FelixKrueger/TrimGalore

Kuleshov MV et al. 2016. Enrichr: a comprehensive gene set enrichment analysis web server 2016 update. Nucleic Acids Res. 44(W1):W90–97. 10.1093/nar/gkw377

Kumsta C, Chang JT, Schmalz J, Hansen M. 2017. Hormetic heat stress and HSF-1 induce autophagy to improve survival and proteostasis in C. elegans. Nat Commun. 8:14337. 10.1038/ncomms14337

Labbadia J et al. 2017. Mitochondrial stress restores the heat shock response and prevents proteostasis collapse during aging. Cell Rep. 21(6):1481–1494. 10.1016/j.celrep.2017.10.038

Leek JT et al. 2012. The sva package for removing batch effects and other unwanted variation in high-throughput experiments. Bioinformatics. 28(6):882–883. 10.1093/bioinformatics/bts034

Liao Y, Smyth GK, Shi W. 2019. The R package Rsubread is easier, faster, cheaper and better for alignment and quantification of RNA sequencing reads. Nucleic Acids Res. 47(8):e47. 10.1093/nar/gkz114

Link CD, Cypser JR, Johnson CJ, Johnson TE. 1999. Direct observation of stress response in Caenorhabditis elegans using a reporter transgene. Cell Stress Chaperones. 4(4):235–242. 10.1379/1466-1268(1999)004%253C0235:doosri%253E2.3.co;2

Liu Y, Chang A. 2008. Heat shock response relieves ER stress. EMBO J. 27(7):1049–1059. 10.1038/emboj.2008.42

Love MI, Huber W, Anders S. 2014. Moderated estimation of fold change and dispersion for RNA-seq data with DESeq2. Genome Biology. 15(12):550. 10.1186/s13059-014-0550-8

Lu H, Koju N, Sheng R. 2024. Mammalian integrated stress responses in stressed organelles and their functions. Acta Pharmacol Sin. 45(6):1095–1114. 10.1038/s41401-023-01225-0

Ma Y, Hendershot LM. 2003. Delineation of a negative feedback regulatory loop that controls protein translation during endoplasmic reticulum stress. J Biol Chem. 278(37):34864–34873. 10.1074/jbc.M301107200

Metcalf MG et al. 2024. Cell non-autonomous control of autophagy and metabolism by glial cells. iScience. 27(4):109354. 10.1016/j.isci.2024.109354

Morley JF, Morimoto RI. 2004. Regulation of longevity in Caenorhabditis elegans by heat shock factor and molecular chaperones. Mol Biol Cell. 15(2):657–664. 10.1091/mbc.E03-07-0532

Neef DW et al. 2014. A direct regulatory interaction between chaperonin TRiC and stress-responsive transcription factor HSF1. Cell reports. 9(3):955–66. 10.1016/j.celrep.2014.09.056

Ryoo HD. 2024. The integrated stress response in metabolic adaptation. J Biol Chem. 300(4):107151. 10.1016/j.jbc.2024.107151

Schindelin J et al. 2012. Fiji: an open-source platform for biological-image analysis. Nat Methods. 9(7):676–682. 10.1038/nmeth.2019

Schröder M, Kaufman RJ. 2005. The mammalian unfolded protein response. Annu Rev Biochem. 74:739–789. 10.1146/annurev.biochem.73.011303.074134

Shi Y, Mosser DD, Morimoto RI. 1998. Molecular chaperones as HSF1-specific transcriptional repressors. Genes & development. 12(5):654–66. 10.1101/gad.12.5.654

Sural S et al. HSB-1/HSF-1 pathway modulates histone H4 in mitochondria to control mtDNA transcription and longevity. Science Advances. 6(43):eaaz4452. 10.1126/sciadv.aaz4452

Taylor RC, Dillin A. 2013. XBP-1 is a cell-nonautonomous regulator of stress resistance and longevity. Cell. 153(7):1435–1447. 10.1016/j.cell.2013.05.042

Torres TC et al. 2022. Surveying Low-Cost Methods to Measure Lifespan and Healthspan in Caenorhabditis elegans. JoVE (Journal of Visualized Experiments). (183):e64091. 10.3791/64091

Walter P, Ron D. 2011. The unfolded protein response: from stress pathway to homeostatic regulation. Science. 334(6059):1081–1086. 10.1126/science.1209038

Wang Y et al. 2024. Metabolic regulation of misfolded protein import into mitochondria Mapa K, Ron D, editors. eLife. 12:RP87518. 10.7554/eLife.87518

Watterson A et al. 2022. Loss of heat shock factor initiates intracellular lipid surveillance by actin destabilization. Cell Rep. 41(3):111493. 10.1016/j.celrep.2022.111493

Wek RC, Anthony TG, Staschke KA. 2023. Surviving and Adapting to Stress: Translational Control and the Integrated Stress Response. Antioxid Redox Signal. 39(4–6):351–373. 10.1089/ars.2022.0123

Williams KW et al. 2014. Xbp1s in Pomc neurons connects ER stress with energy balance and glucose homeostasis. Cell Metab. 20(3):471–482. 10.1016/j.cmet.2014.06.002

Ye J et al. 2000. ER stress induces cleavage of membrane-bound ATF6 by the same proteases that process SREBPs. Mol Cell. 6(6):1355–1364. 10.1016/s1097-2765(00)00133-7

Zevian SC, Yanowitz JL. 2014. Methodological Considerations for Heat Shock of the Nematode Caenorhabditis elegans. Methods. 68(3):450–457. 10.1016/j.ymeth.2014.04.015

Zha J, Ying M, Alexander-Floyd J, Gidalevitz T. 2019. HSP-4/BiP expression in secretory cells is regulated by a developmental program and not by the unfolded protein response. PLOS Biology. 17(3):e3000196. 10.1371/journal.pbio.3000196

